# Selective convergence and graded divergence of hippocampal and amygdala subregions using functional connectivity

**DOI:** 10.64898/2026.07.06.736898

**Authors:** Doruk Yiğit Erigüç, Mylla Marsiglia, Alexandra John, Şeyma Bayrak, Bin Wan, Anton Jakovčić, Jordan DeKraker, Jessica Royer, Boris C. Bernhardt, Sofie L. Valk

## Abstract

The hippocampus and amygdala are neighboring medial temporal lobe structures linked to memory and affect, yet how their subregions are jointly embedded within distributed isocortical systems remains unclear. Using resting-state fMRI from 722 Human Connectome Project Young Adult participants, we mapped hippocampal and amygdalar subregions within a unified cortex-wide framework, quantifying subregion-to-cortex connectivity via Pearson correlation (broad co-fluctuation) and GLASSO partial correlation (relatively more direct functional association). We introduced two count-based metrics: dominance (relative hippocampal vs. amygdalar representation) and sharedness (balanced co-representation). Direct associations showed both structures sharing coupling with paralimbic areas and, more modestly, default mode regions, while broader co-fluctuations extended into somatomotor and paralimbic networks. Divergence patterns depended on the estimator: hippocampal subregions preferentially coupled with default-mode and visual networks under direct association, while amygdalar nuclei favored ventral attention and limbic networks; broader co-fluctuations additionally implicated somatomotor cortex for amygdala and visual cortex for hippocampus. These principles held at the subfield/nucleus level, varying along the hippocampal long axis and identifying the paralaminar nucleus as the most hippocampus-like amygdalar subregion. Data-driven connectivity gradients confirmed both systems’ separation and fine-scale interdigitation. Hippocampal and amygdalar subregions are thus embedded in cortex not as discrete systems, but through structured, spatially organized co-representation.

## Introduction

The hippocampus and amygdala are neighboring structures in the medial temporal lobe (MTL) (Kiernan, 2012). Both are embedded within distributed cerebral cortical networks, but show partly distinct large-scale connectivity profiles. Hippocampal connectivity is often described in terms of large scale cortico-hippocampal memory systems. This structure interacts with the cerebral cortex through entorhinal, perirhinal, and parahippocampal interfaces and shows distinct coupling with anterior-temporal/medial-prefrontal and posterior-medial/parietal networks along its extent (Ranganath & Ritchey, 2012; Ritchey et al., 2015; Vos de Wael et al., 2018; Paquola et al., 2020; Barnett et al., 2021; Genon et al., 2021; Huang et al., 2021). The amygdala shows especially prominent connections with medial and orbitofrontal prefrontal cortex alongside broader temporal, paralimbic, and salience-related cortical coupling (Ghashghaei et al., 2007; Bickart et al., 2014; Barbas & García-Cabezas, 2017; Elvira et al., 2022; Klein-Flügge et al., 2022; Auer et al., 2025). However, this distinction is not absolute: anatomical studies in non-human primates show that amygdala and hippocampal formation efferents converge in medial temporal cortices, including entorhinal, prorhinal, and perirhinal regions, and that both structures show complementary projections to medial and orbital prefrontal territories (Saunders & Rosene, 1988a; Saunders et al., 1988b; Aggleton et al., 2015). These partially overlapping cortical areas provide a circuit-level rationale for studying the hippocampus and amygdala within a shared cortical reference frame. Yet most human studies have characterized the hippocampus and amygdala separately, often at the level of whole structures or preselected cortical targets. Because both structures are internally heterogeneous, it remains unclear how hippocampal subfields and amygdala subnuclei are jointly organized with respect to the cortex. Importantly, shared cortical embedding can have different meanings. A cortical parcel may appear shared because it receives relatively direct or conditionally specific input from both structures, because both structures participate in the same broader network-level co-fluctuation pattern, or because their apparent association is mediated by other areas in the brain. Distinguishing these possibilities is essential for translating animal circuit findings on hippocampus-amygdala pathways and cortical convergence to human fMRI.

Both structures are implicated in a broad range of distinct but also shared functions. The hippocampus has classically been associated with spatial navigation (Tolman, 1948; O’Keefe & Nadel, 1978) and episodic memory (Scoville & Milner, 1957) but contemporary perspectives view it as supporting domain general relational processing (Eichenbaum et al., 1999; Eichenbaum, 2017). It represents the relational structure of experiences, encoding multidimensional associations among objects, locations, entities, rules and events (often referred to as cognitive maps; Eichenbaum et al., 1999; Eichenbaum, 2017; Behrens et al., 2018; Bellmund et al., 2018; Whittington et al., 2020, 2022; Courellis et al., 2024) and can be divided along its longitudinal axis on both functional, structural and molecular bases (Fanselow & Dong, 2010; Poppenk et al., 2013; Strange et al., 2014; Vogel et al., 2020; Genon et al., 2021; Dalton et al., 2022; Nordin et al., 2025) or along the transverse (proximodistal) axis to its canonical subfields based on cytoarchitecture or function (Insausti & Amaral, 2004; Amunts et al., 2005; Genon et al., 2021). Likewise, the amygdala has historically been considered to be the fear center of the brain based on lesion and fear conditioning studies (Klüver & Bucy, 1939; Weiskrantz, 1956; LeDoux, 2000, 2003). Recent work, however, has viewed the amygdala as a social-affective hub (Bickart et al., 2014) with multiple functions supported by its different subnuclei such as valence (also positive; Jin et al., 2015) and salience encoding (Kong & Zweifel, 2021), regulation of social and reproductive behaviors (Ignacio et al., 2025), olfactory processing and integrating sensory information to build stimulus-outcome associations (Wassum & Izquierdo, 2015). Despite their distinct roles, the hippocampus and amygdala work synergistically in a range of brain functions including encoding/retrieval of emotionally salient memories (Fastenrath et al., 2014; Hermans et al., 2014;

Qasim et al., 2023; Costa et al., 2022, 2025) and working memory (Li et al., 2023). Emotional arousal signals from the amygdala can enhance or modulate hippocampal memory formation, while contextual information from the hippocampus can regulate amygdala responses (Kim & Cho, 2020; Qasim et al., 2023; Costa et al., 2022, 2025). This interplay is likely supported by direct bidirectional connections between the ventral hippocampus (especially CA1 and subiculum) and the basolateral amygdala, as well as indirect influences via parallel targeting of cortical areas such as medial temporal and orbital prefrontal cortices (Saunders & Rosene, 1988a; Saunders et al., 1988b; Pitkänen et al., 2000; Ghashghaei et al., 2007; Aggleton et al., 2015; McDonald & Mott, 2016).

The interconnectedness of hippocampal subfields and amygdalar nuclei may have evolutionary origins, rooted in the dual-origin theory of cortical organization. This theory proposes that the mammalian cortex expanded from two ancient allocortical anchors: a hippocampal/archicortical origin and an olfactory-piriform/paleocortical origin closely related to amygdalar and paralimbic territories (Sanides, 1962, 1970; Pandya & Yeterian, 1985; García-Cabezas et al., 2019, 2023). From this perspective, hippocampus and amygdala/piriform cortex may not simply connect to isocortex as two independent medial temporal structures, but may reflect partially distinct allocortical roots whose influence is expressed along parahippocampal and paraolfactory cortical gradients. Prior work suggests that hippocampal cortical coupling varies along the anterior-posterior axis, with posterior portions more closely linked to default mode systems and anterior portions more closely linked to limbic and attention-related systems (Ranganath & Ritchey, 2012; Ritchey et al., 2015; Fritch et al., 2021; Panitz et al., 2021), though findings are somewhat conflicting (Vos de Wael et al., 2018; Katsumi et al., 2023). The amygdala is connected to limbic, default mode, somatomotor, salience/ventral attention and temporoparietal networks (Bickart et al., 2014; Kerestes et al., 2017) however, amygdala-subnuclei-specific mapping to cortical networks remains inconsistent (Bzdok et al., 2013; Kerestes et al., 2017; Sylvester et al., 2020; Elvira et al., 2022; Klein-Flügge et al., 2022; Auer et al., 2025). Critically, animal anatomical studies suggest that hippocampus and amygdala relate to cortex not only through broad systems-level co-engagement, but through a combination of direct projections, indirect polysynaptic influences, and partial convergence onto shared cortical targets, particularly in medial and orbitofrontal territories (Saunders & Rosene, 1988a; Saunders et al., 1988b; Ghashghaei et al., 2007; Aggleton et al., 2015). However, most human functional MRI studies rely on pairwise correlation, which cannot determine whether apparent similarity in cortical connectivity profiles reflects direct coupling, indirect network-mediated dependence, or both. Given their shared roles in memory, emotion and attention, and their known interconnectivity and overlapping cortical targets, studying the hippocampus and amygdala together is essential.

Here, we aimed to characterize how hippocampal and amygdalar subregions are jointly organized in reference to the cerebral cortex (isocortex), leveraging the Human Connectome Project Young Adult (HCP-YA) dataset. First, we sought to identify where the two structures show distinct versus convergent functional connectivity (FC) profiles in the isocortex. For this purpose, we derived explicit cortical mapping metrics to quantify predefined aspects of divergence and convergence in a more interpretable way, including hippocampal versus amygdalar dominance and sharedness. Next, we asked whether these patterns can be summarised by continuous, low-dimensional axes of joint organization. For the second aim we used joint hippocampus-to-cortex and amygdala-to-cortex FC gradient mapping to derive low dimensional axes of subregion-to-cortex connectivity without prespecifying organizational features (Coifman et al., 2005; Margulies et al., 2016; Vos de Wael et al., 2018). Placing hippocampus and amygdala subregions in a shared embedding space enabled us to compare their separation, overlap and graded variation in their cortical connectivity profiles. Importantly, our use of both direct and indirect functional connectivity markers captured different aspects of isocortical dependence across regions. Pearson correlation captures dense co-fluctuation structure that includes both direct and indirect connections, whereas graphical lasso (GLASSO) based partial correlation provides a sparser, conditionally specific estimate intended to reduce indirect effects, allowing us to capture relatively more direct connections (Friedman et al., 2008; Smith et al., 2011; Liégeois et al., 2020; Peterson et al., 2025). We hypothesized that hippocampal and amygdalar subregions would exhibit both structure specific cortical preference zones and regions of convergence and that joint gradients would also recapitulate these features, showing separation in some axes and shared functional projections in others. We expected Pearson correlation to reveal broader shared co-fluctuation, whereas GLASSO would emphasize more focal conditional associations that are closer to anatomical findings.

## Methods

### Participants

We leveraged neuroimaging data from the Human Connectome Project Young Adult S-1200 2025 reprocessed release (HCP-YA, Van Essen et al., 2013). Details on data acquisition and preprocessing have been described elsewhere in detail (Glasser et al., 2013; Van Essen et al., 2013), but in the following briefly summarized.

From the original cohort of 1,200 participants, we included 722 subjects (mean age = 28.48 ± 3.75 years; 380 females) who were retained after quality assessment. Participants were excluded based on the following criteria: *i)* subjects with imaging-related issues reported on the official HCP website (https://wiki.humanconnectome.org), *ii)* incomplete acquisition of the four resting-state fMRI (rfMRI scans), *iii)* mean relative root mean square (RMS) displacement exceeding 0.2 mm, *iv)* absence of fMRI image reconstruction using the r227 (Recon2) algorithm, *v)* unavailability of the required imaging data in the HCP release used for analysis, and *vi)* failure of subject-level cortical, hippocampal, or amygdala processing, defined as missing required outputs, incomplete regional parcellation, invalid or zero-variance regional time series, excessive missing values, or invalid connectivity-matrix dimensions.

### MRI Acquisition

Neuroimaging data for the HCP-YA cohort were acquired at a single imaging site at Washington University in St. Louis using a customized Siemens Skyra 3T MRI scanner ("Connectom") equipped with a standard Siemens 32-channel head coil. High-resolution structural T1-weighted (T1W) images were obtained through a three-dimensional magnetization-prepared rapid gradient-echo (3D MPRAGE) sequence (TR = 2.4 s; TE = 2.14 ms; TI = 1000 ms), yielding isotropic voxels with a spatial resolution of 0.7 mm. Resting-state fMRI data were acquired using a blood oxygen level-dependent (BOLD) gradient-echo echo-planar imaging (GE-EPI) sequence at 3 Tesla with an echo time (TE) of 33 ms (Smith et al., 2013). Data acquisition employed a multiband acceleration factor of 8 with blipped-CAIPIRINHA, enabling a high temporal resolution of 0.72 seconds (TR) while maintaining an isotropic spatial resolution of 2 mm. Each participant underwent one hour of rfMRI acquisition, divided into four runs of 15 minutes across two separate sessions. During scanning, participants were instructed to remain still, keep their eyes open, fixate on a white cross, and refrain from engaging in any specific thoughts or falling asleep. To minimize susceptibility-induced signal distortions, two runs were acquired with left-to-right (L-R) phase encoding, while the remaining two runs with right-to-left (R-L) phase encoding.

### MRI Preprocessing

Structural and functional MRI data were processed using the HCP minimal preprocessing pipeline. Structural MRI underwent gradient distortion correction, bias field correction, registration to the MNI152 standard space, and tissue segmentation into gray matter, white matter, and cerebrospinal fluid (CSF), and cortical surface reconstruction. These procedures were implemented using established neuroimaging tools, including FSL (Jenkinson et al., 2012), Freesurfer (Fischl, 2012) and Connectome-Workbench (Marcus et al., 2013). RfMRI images underwent gradient unwarping, motion correction, susceptibility distortion correction using field maps, high-pass temporal filtering (hp2000, effectively linear detrending) to attenuate slow signal drifts, co-registration to the T1W image, spin-echo based intensity bias field correction (SEBASED). Subsequently, data were denoised using ICA-FIX, Reclean and temporal ICA (tICA), and registered to MNI152 space, converted to CIFTI format, and aligned across cortical surfaces using MSM-All.

For subcortical analyses, we performed additional preprocessing steps to achieve optimal alignment between functional and anatomical data within a common subject-specific volumetric space. We resampled the native space T1W image to 2 mm isotropic resolution to match the spatial resolution of the rfMRI data. Then we used the subject native volumetric functional data (transformed using the provided transformation files, using cubic spline interpolation) for amygdala and hippocampal time series extraction. Final hippocampal time series were obtained after additional HippUnfold processing. In parallel, cortical time series were derived from the HCP-cleaned fsLR surface data. Accordingly, cortical, amygdalar and hippocampal BOLD signals were analyzed within the anatomical framework most appropriate for each structure.

### Cortical Parcellation and Subcortical Segmentation

Cortical parcellation was performed using HCP Multi-Modal Parcellation 1.0 (HCP-MMP 1.0, will be referred to as Glasser parcellation; Glasser et al., 2016) applied to the fsLR 32k cortical surfaces derived from MSMAll-aligned CIFTI data. This parcellation divides the cortex into 360 areas. We excluded two hippocampus-assigned parcels from the Glasser parcellation because we already included hippocampus subregions obtained using HippUnfold, leaving 358 cortical parcels for analysis.

We segmented the amygdala into nine subnuclei per hemisphere in each participant’s native space **(Fig. 1A)** using the segment_subregions module implemented in FreeSurfer, following the probabilistic atlas approach described by Saygin et al. (2017). This procedure generated amygdala subnuclei label maps at 0.333 mm isotropic resolution. Rather than directly downsampling these labels into hard masks, we constructed mass-conserving soft masks at 2 mm isotropic resolution to match the functional data. Hard masks assign each voxel entirely to one nucleus, whereas soft masks preserve fractional occupancy when high-resolution labels are pooled to 2 mm, reducing partial-volume loss. Specifically, high resolution binary label maps for each hemisphere and nucleus were first resampled using nearest-neighbor interpolation onto a 0.333 mm grid aligned to the 2 mm T1W reference image and subsequently pooled exactly (6 x 6 x 6) to generate voxel-wise fractional masks at 2 mm resolution.

**Figure 1.**
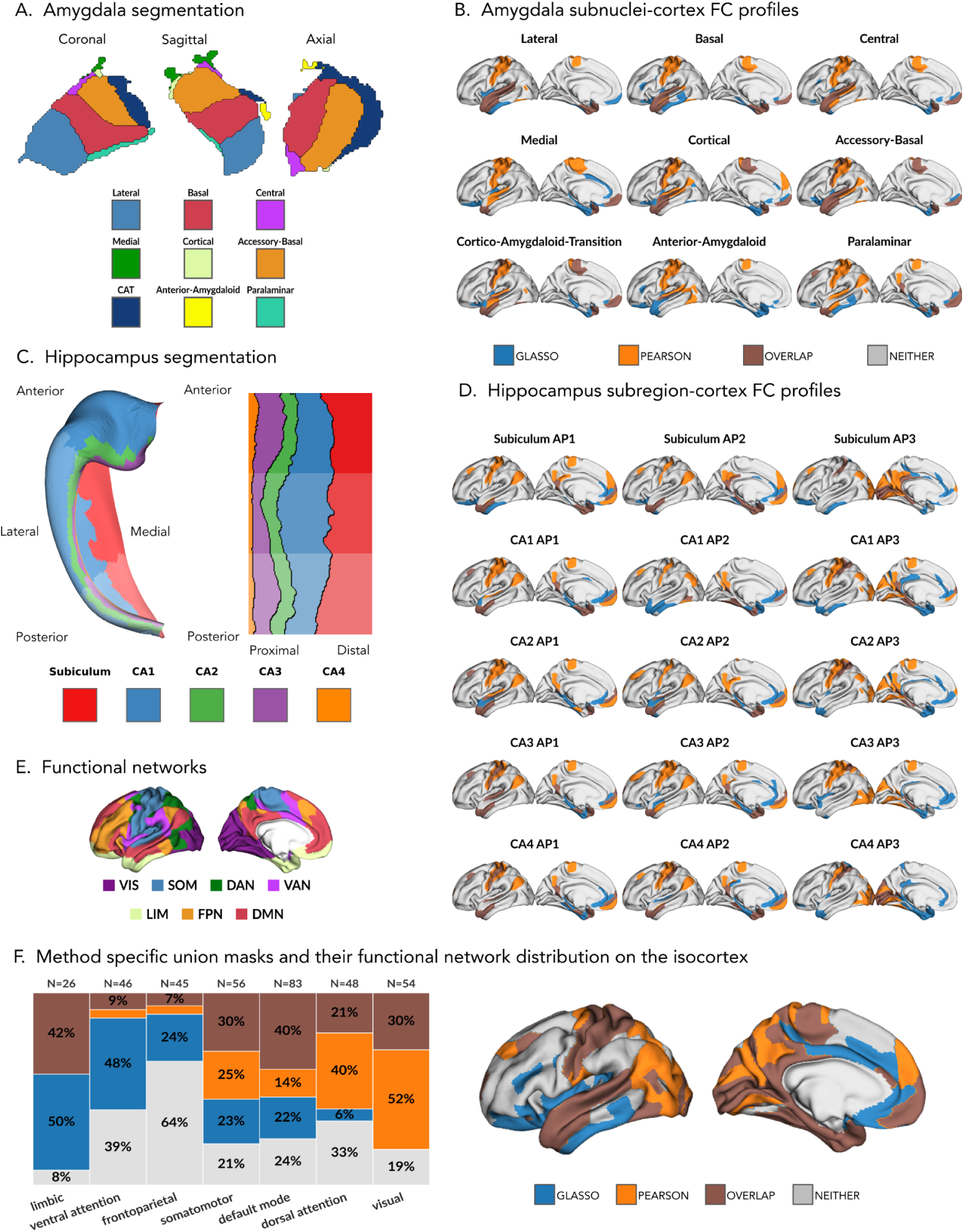
Amygdalar and hippocampal subregion delineations and subregion-to-cortex functional connectivity profiles. For each estimator, union of seed-cortex FC maps were used as masks for the following dominance and sharedness analyses. For each seed, cortical maps show the top-10% positive seed-cortex connections at the group level, projected from Glasser parcels to the fsLR32k cortical surface (only left hemisphere shown here). For GLASSO, top-10% edges are defined by support prevalence across subjects and are shown in blue; for Pearson, top-10% edges are defined by connection strength (positive correlations) and are shown in orange. Their conjunction is shown in brown. Glasser parcels for the hippocampus were excluded. **A)** Amygdala segmentation into nine subnuclei, illustrated in volumetric representations (Saygin et al., 2017). **B)** Amygdalar subnuclei-to-cortex FC profiles, each tile shows the cortical FC of a single amygdala subnucleus’ most prevalent or strongest connections. **C)** Hippocampus segmented into five subfields and further subdivided into three anterior-posterior divisions (anterior, middle, and posterior), illustrated in both folded and unfolded surface representations (DeKraker et al., 2022). **D)** Hippocampal subregion-to-cortex FC profiles shown separately for the anterior (AP1), middle (AP2), and posterior (AP3) portions of each subfield. Each tile shows the cortical FC of a single hippocampal subregion’s most prevalent or strongest connections. Rows correspond to subfields and columns to A-P bins. **E)** Parcels belonging to different Yeo-7 functional networks shown on the cortical surface (visual network = VIS, somatomotor network = SOM, dorsal attention network = DAN, ventral attention network = VAN, limbic network = LIM, frontoparietal network = FPN, default mode network = DMN). **F)** GLASSO and Pearson union masks, their overlap, and cortical regions not covered by either mask are displayed on the cortex (left). The distribution of individual masks (y-axis) across the Yeo-7 functional networks (x-axis) is summarized in a stacked bar plot showing the proportional overlap of each mask within each network (right).

We used HippUnfold 1.5.1 (DeKraker et al., 2022) to define hippocampal subfield labels (CA1, CA2, CA3, CA4 and subiculum) and corresponding subject-specific hippocampal surfaces, based on the native-space T1W image. HippUnfold applies a U-Net-based deep convolutional neural network architecture (Isensee et al., 2021) to generate detailed segmentations of hippocampal gray matter. The resulting segmentations and unfolded representations have been extensively validated against histological reference data (DeKraker et al., 2022, 2023, 2025). Functional data were sampled onto these surfaces and summarized at the parcel level. To further characterise longitudinal organization, we divided each hippocampal subfield into three equal sections along the unfolded longitudinal axis, yielding 15 hippocampal parcels per hemisphere for each participant in their native space (**Fig. 1C**).

Across all, the final node set comprised 18 amygdala subnuclei, 30 hippocampal parcels, and 358 cortical parcels, yielding a total of 406 regions.

### Regional time series extraction

Cortical parcel time series were obtained from the HCP-cleaned fsLR surface data by averaging vertex-wise time series within each retained Glasser parcel. Amygdala nucleus time series were extracted in subject-specific native 2 mm volumetric space using fractional mask-weighted averaging of the concatenated 4D BOLD data. For each nucleus, voxel-wise BOLD signals were weighted according to the corresponding fractional occupancy values derived from soft masks.

Hippocampal parcel time series were extracted from HippUnfold derived hippocampal functional surfaces by averaging functional values within each parcel.

### Data Analysis

#### Functional Connectivity Estimation

For each participant, regional time series from the 406-node set (358 cortical parcels, 18 amygdala nuclei and 30 hippocampal parcels) were concatenated across the four rfMRI runs. Each node time series was then z-scored across the concatenated time axis within the participant, such that each regional signal had mean zero and unit variance before FC estimation. Pairwise Pearson correlation coefficients were then computed between all node pairs to obtain a 406 × 406 subject-level FC matrix.

Additionally, we estimated sparse partial correlation FC matrices using the graphical lasso (GLASSO) implementation provided in the Brain Activity Flow (Actflow) Toolbox (Cole et al., 2016; Peterson et al., 2025). For each subject, parcel-wise time series were z-scored and used to compute an empirical covariance matrix. The L1 regularization parameter (λ_1_) was optimized individually for each subject using 4-fold blocked cross-validation, with each fold corresponding to one rfMRI run. The optimal λ value was selected from a default logarithmically spaced parameter grid by minimizing the mean negative log-likelihood on held-out folds. The final GLASSO model was then refitted on all time points using the selected λ value to obtain a subject-specific sparse partial correlation matrix.

Subject-specific GLASSO regularization parameters selected through blocked cross-validation varied across participants (mean λ = 0.020, SD = 0.004). As expected, larger λ values were associated with sparser subject-level graphs (r = -0.96 for log10 λ and edge density). Mean relative RMS displacement showed weak associations with λ and density (r = 0.27 and r = -0.22, respectively), indicating that head motion exerted only a minor influence on matrix sparsity and did not trivially determine the GLASSO solutions.

Subject-level Pearson correlation and GLASSO matrices were Fisher z-transformed prior to group-level averaging. The resulting group-level Pearson and GLASSO z-matrices were then inverse z-transformed to obtain the final group-level FC matrix used in downstream analyses. During GLASSO group averaging, zero valued edges were treated as missing values rather than true correlations, as zeros reflected absent edges imposed by regularization and would otherwise bias the group mean toward zero.

We additionally quantified GLASSO support prevalence as the proportion of participants for whom a given subregion-cortex partial-correlation edge was positive.

### Method Specific Masks

To characterize the spatial distribution of hippocampal and amygdalar connectivity with the cortex, we extracted a 48 x 358 submatrix corresponding to subregion-to-cortex connections. We generated method-specific union masks to isolate robust connectivity patterns for downstream analyses. For the Pearson correlation matrices, we restricted the analysis to the union of the top 10% strongest positive-signed connections for each subregion (hereafter referred to as Pearson mask). For the GLASSO-derived partial correlation matrices, we restricted the analysis to the union of the top 10% most prevalent positive connections for each subregion, where prevalence was defined as the consistency of positive edges across the subject-specific sparse connectivity matrices (hereafter referred to as GLASSO mask). These masks were then used for seed-level cortical connectivity visualizations and for all count-based analyses, including dominance, sharedness, seed-specific preference scores, and strength-based corroboration. They were not used to restrict the joint gradient estimation. For gradient analyses, the full retained subregion-to-cortex matrix was used, with the exception that GLASSO matrices were additionally consensus-masked as described below.

### Dominance and sharedness metrics

For estimates derived from both methods, we generated two complementary count-based cortical mapping metrics within the corresponding method-specific masks: dominance and sharedness.

First, we computed dominance indices to quantify the relative contribution of hippocampal versus amygdalar connectivity to each cortical parcel. For every cortical parcel, we counted the number of suprathreshold seed-to-cortex connections (connections included within the respective method-specific mask) originating from each seed group and normalized these counts by the total number of seeds within the corresponding structure (hippocampus or amygdala). Dominance was then defined as a signed normalized contrast ranging from -1 to +1, where -1 indicated exclusive hippocampal contribution, +1 indicated exclusive amygdala contribution and 0 reflected equal normalized contributions from both structures. Cortical parcels with no suprathreshold connections from either seed group were excluded.

Second, we computed a sharedness index quantifying the degree of convergent hippocampal and amygdalar connectivity at each cortical parcel. This metric incorporated both the overall magnitude of joint targeting (defined as the sum of normalized hippocampal and amygdalar contributions) and the balance between these contributions (defined as the ratio of the smaller to the larger normalized contribution). Consequently, sharedness was maximal when both hippocampal and amygdalar inputs were simultaneously strong and proportionally balanced. The resulting sharedness metric ranged from 0, indicating either absent shared targeting or complete imbalance, to 2, indicating maximal and perfectly balanced co-representation. Distributions of dominance and sharedness values were subsequently assessed and visualized across Yeo-7 cortical networks (Yeo et al., 2011).

To evaluate the robustness of these metrics with respect to threshold selection of top 10%, dominance and sharedness maps were additionally recomputed using top 5% and top 15% masks for both Pearson and GLASSO FC estimates. Pairwise similarity between these alternative threshold maps and the primary top 10% maps used in the main analyses were then quantified.

### Strength-based corroboration of dominance

For evaluating whether the count-based dominance metric reflected underlying differences in connectivity magnitude, we computed a complementary strength-based dominance measure. For each cortical parcel, positive amygdala-to-cortex and hippocampus-to-cortex connectivity values were separately averaged within the respective method-specific support. We then derived parcel-wise amygdala-minus-hippocampus contrast maps based on these strength estimates and assessed their correspondence with the count-based dominance metric using parcel-wise correlation analyses. Crucially, this correlational analysis was restricted to parcels exhibiting at least one suprathreshold connection from both amygdala and hippocampus, thereby avoiding inflation driven by extreme dominance values (+-1 arising from unisource connectivity).

### Subregion-seed-specific cortical preference

To quantify how individual amygdala and hippocampus subregions preferentially contributed to cortical patterns of dominance or sharedness, we computed seed-specific preference indices using a leave-one-seed-out (LOSO) procedure. For each seed, group-level dominance and sharedness maps were recalculated after excluding that seed’s contribution, providing seed-independent reference maps and reducing circularity. Each seed’s cortical connectivity profile within its own top-10% method-specific mask was then used to weight these LOSO-derived maps by applying connectivity-weighted averaging, yielding seed-specific preference scores for dominance and sharedness.

For dominance, positive values indicated preferential weighting toward amygdala-dominant cortical regions, negative values indicated preferential weighting toward hippocampus-dominant regions, and values near zero reflected balanced or weak preference. For sharedness, higher values indicated preferential weighting toward cortical regions receiving strong and balanced input from both structures.

### Amygdala-hippocampus seed-to-seed coupling

To characterize intrinsic cross-structure interactions, we extracted the 18 x 30 amygdala-hippocampus block from the full connectivity matrix. Group-level positive coupling was summarized as the unconditional positive mean across subjects. For visualization purposes, both raw coupling matrices and column-wise min-max normalized matrices were examined, with the latter emphasizing relative interaction structure independent of absolute magnitude. Ipsilateral blocks were highlighted in the main figure for visual clarity.

We further tested whether each seed’s intrinsic cross-structure coupling strength was related to its extrinsic cortical sharedness preference. Specifically, seed-wise intrinsic amygdala-hippocampus coupling strength was correlated with the corresponding LOSO-derived sharedness preference score using Spearman correlation. In addition, partial Spearman correlations were computed while controlling for each seed’s total positive cortical connectivity strength. Resulting *p-*values were corrected for multiple comparisons using false discovery rate (FDR) correction.

### Joint amygdala-hippocampus-to-cortex gradients

For functional gradient estimation, we used the unmasked 48 x 358 amygdala-hippocampus-to-cortex group-average FC matrices from both FC estimates as input. For the GLASSO group average FC matrix, we additionally applied a consensus mask to reduce unstable edges by retaining only seed-to-cortex connections that were present as positive edges in at least 10% of subjects. Gradient estimation was performed separately for the left and right hemisphere seed sets to bilateral cortex to prevent the dominant gradient structure from being driven by global left-right differences between seed hemispheres, using the BrainSpace toolbox (Vos de Wael et al., 2020). We first computed an affinity matrix (24 x 24 for each hemisphere) that captured the similarity of cortical connectivity profiles between each pair of seed regions, using the normalized-angle kernel. This affinity matrix was then thresholded row-wise (retaining the top 10% strongest similarities) to create a sparse graph, emphasizing dominant connectivity relationships while reducing potential noise. Diffusion embedding was then applied to this sparse graph to identify low-dimensional principal axes of variation, referred to as gradients (Coifman et al., 2005; Margulies et al., 2016). The resulting gradients represent the main organizational principles describing how these hippocampal and amygdalar subregions functionally relate to the cortex. Gradients obtained from the left and right hemisphere were subsequently aligned, with the left hemisphere being the template, using Procrustes alignment to facilitate comparability across hemispheres. We focused on the first two gradients for each estimator.

To aid spatial interpretation, gradient loadings were projected back to the cortex by correlating each seed-gradient vector (48 seed scores) with each cortical parcel’s 48 length seed-to-subregion connectivity vector. This projection was performed across the full retained cortical parcellation, after excluding the two hippocampal Glasser parcels. For GLASSO, the projection used the same consensus-masked group-level connectivity matrix used for gradient estimation. Thus, the cortical projection maps summarize how each joint amygdala-hippocampus gradient relates to cortex-wide subregion-to-cortex connectivity profiles on the fsLR 32k surface, rather than only to parcels included in the top-10% method-specific masks. The distribution of cortical projection values across Yeo-7 networks ( Yeo et al., 2011) was assessed and visualized.

To relate our cortical projection maps to established axes of macroscale cortical organization, we quantified their correspondence with the first three canonical FC gradients (derived from our sample using pairwise Pearson correlations) using an approach analogous with Margulies et al. (2016). Canonical gradients were obtained in fsLR-32k space and summarized at the Glasser-parcel level to match our cortical representation. We computed parcelwise correlations between each projected joint gradient map and the first three canonical FC cortical gradients summarized in Glasser parcel space. We also quantified correspondence between projected joint gradients and the cortex-level amygdala-hippocampus dominance and sharedness maps.

### Spatial statistics and multiple-comparison correction

We assessed spatial correspondence between cortical parcelwise maps using parcel-based Moran spectral randomization using Neuromaps (Wagner & Dray, 2015; Markello et al., 2022), which preserves cortical spatial autocorrelation under the null. Unless otherwise noted, significance for cortex-level map-to-map correspondence was assessed using 1000 spatial null permutations generated with Moran spectral randomization. Seed-level association analyses and correlations between cortical maps were corrected for multiple comparisons using FDR.

## Results

### Shared and estimator-specific cortical coupling of amygdala and hippocampus subregions (Figure 1)

To initially characterize the cortical organisation of amygdalar subnuclei (9 subregions per hemisphere) and hippocampal subfields (15 subregions per hemisphere), we examined the seed-wise FC profiles of all delineated amygdala and hippocampus subregions with the cortex. These FC profiles were subsequently evaluated within the method-specific masks derived from GLASSO partial correlation and Pearson correlation analyses (**Fig. 1B, D**). Both connectivity estimators revealed structured yet heterogeneous seed-to-cortex connectivity patterns; however, they emphasized partially distinct cortical territories. GLASSO-derived connectivity profiles were more spatially focal and subregion-specific, with connectivity concentrated predominantly within limbic and paralimbic territories, particularly medial and orbitofrontal, and cingulate (especially for hippocampal subregions) cortices. In contrast, using Pearson correlation, the majority of amygdalar nuclei exhibited overlapping connectivity patterns within somatomotor cortex, superior temporal cortex, and medial prefrontal regions. Hippocampal parcels demonstrated pronounced variation along the anterior-posterior axis, accompanied by additional graded differences across subfields. Overall, the two approaches captured a broadly similar organizational architecture while emphasizing differential aspects of FC. The GLASSO approach emphasized spatial specificity of positive functional coupling, whereas the Pearson approach highlighted broader patterns of positive co-fluctuation.

Comparison of the method-specific union masks across the subregion seeds revealed both overlapping and clear estimator-specific territories. The GLASSO-derived mask showed a sparser and more spatially focal pattern, whereas the Pearson-derived mask exhibited broader overall cortical coverage. The network-level distribution of these masks was markedly non-uniform. The default mode network showed a high degree of overlap between the two masks, whereas limbic and ventral attention networks were more strongly represented in GLASSO-specific components. In contrast, dorsal attention and visual networks were preferentially represented in the Pearson-specific component. The frontoparietal network was largely absent from both masks, while the somatomotor network displayed a mixed pattern of coverage across estimators.

### Dominance and sharedness reveal complementary cortical organization (Figure 2A-B)

To move beyond descriptive overlap and quantify how amygdalar and hippocampal systems jointly target the cortex, we computed parcel-wise dominance and sharedness metrics within each method-specific cortical mask (**Fig. 2A-B**). These two metrics capture complementary aspects of cortical organization: dominance reflects the relative preference for amygdalar- versus hippocampal-cortical FC, whereas sharedness identifies cortical regions that are jointly targeted by both systems in a relatively balanced manner.

**Figure 2.**
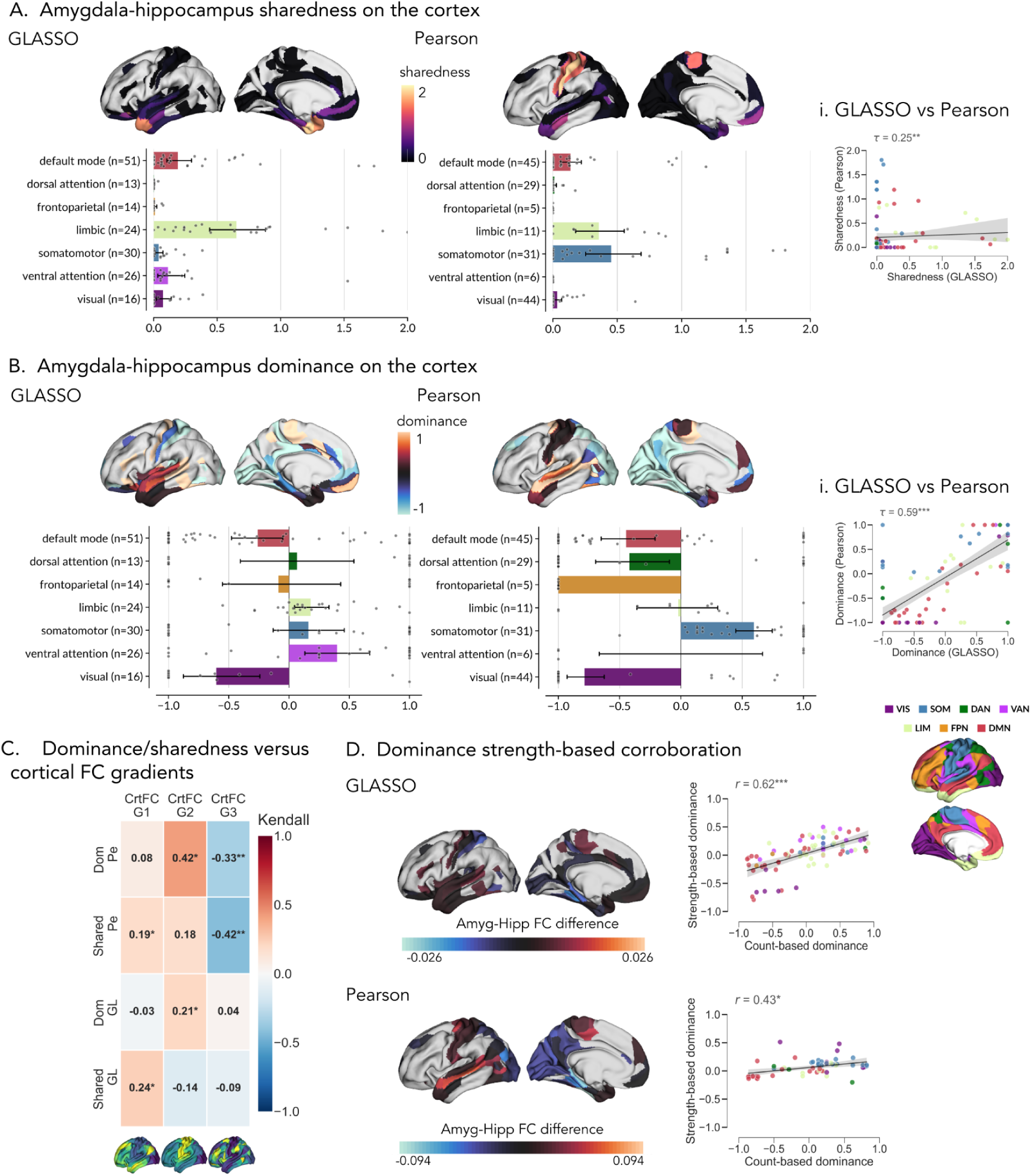
Dominance/sharedness model reveals complementary hippocampus-amygdala organization across cortex. Only left hemisphere brain maps are shown for visualization purposes, analyses are conducted on the entire brain. Right hemisphere results can be found on **Supplementary Figure 3**. For each cortical parcel, we quantified (i) dominance: relative preference for amygdala vs hippocampus connectivity, and (ii) sharedness: balanced co-representation of both structures. The model is computed from seed-wise Top-10% parcel memberships (after structure-wise normalization), such that dominance reflects the signed difference between the number of amygdala and hippocampus contributions (positive = amygdala-dominant, negative = hippocampus-dominant), whereas sharedness is high only where both contributions are simultaneously strong (high count) and balanced. All cortical maps are accompanied by Yeo-7 summaries (bars = network means, lines = 95% CI, dots = parcels). Asterisks denote spatial correspondence significance using Moran spectral randomization (* *p_MSR_* < 0.05, ** *p_MSR_* < 0.01, *** *p_MSR_* < 0.001). **A)** Cortical sharedness maps for GLASSO and Pearson. Higher values indicate stronger balanced hippocampus-amygdala co-representation. Cross-estimator correspondence scatter plots (**i**) comparing GLASSO vs Pearson sharedness. **B)** Cortical dominance maps for GLASSO and Pearson. Positive values indicate amygdala preference; negative values indicate hippocampal preference; values near zero indicate relatively balanced contributions. Cross-estimator correspondence scatter plots (**i**) comparing GLASSO vs Pearson dominance. **C)** Heatmaps depict parcel-wise correlations between GLASSO (GL) and Pearson (Pe) derived dominance (Dom) and sharedness (Shared) maps and the first three cortical functional gradients (CrtFC G1, G2, and G3; derived using Pearson). Asterisks indicate the significance of spatial correspondence assessed via Moran Spectral Randomization and FDR-corrected (* *q_MSR_*<0.05, \*\**q_MSR<_*0.01, *** *q_MSR_*<0.001). **D)** Strength-based validation of the count-based model: FC difference maps (amygdala-cortex FC - hippocampus-cortex FC) for GLASSO (left) and Pearson (right) approaches. Scatter plots display FC-difference-based dominance against count-based dominance for GLASSO (left) and Pearson (right) approaches.

Sharedness showed a markedly focal distribution in the GLASSO estimator, with highest values concentrated in perirhinal, entorhinal, orbitofrontal, piriform, temporopolar, anterior insular and subgenual cortices. Sharedness was predominantly expressed within the limbic network, with more modest contributions from the default mode and ventral attention networks and near-zero values across the remaining networks. In contrast, the Pearson estimator produced a broader distribution of sharedness, with the strongest effects observed in the somatomotor network and secondary contributions from the limbic network. The default mode network showed modest sharedness, whereas sharedness for the remaining networks was minimal. Overall, balanced amygdalar-hippocampal co-representation was restricted to a limited subset of cortical regions and was strongly estimator-dependent, with GLASSO emphasizing a focal paralimbic/limbic core and Pearson revealing a broader pattern.

Dominance maps revealed a complementary organization reflecting amygdalar-weighted versus hippocampal-weighted cortical territories. In the GLASSO solution, the visual network was clearly hippocampus-dominant, ventral attention network showed the strongest amygdala dominance, and limbic and somatomotor networks exhibited mild amygdala preference. The default mode and frontoparietal regions were slightly hippocampus-biased, whereas the dorsal attention network was near neutral. In the Pearson solution, the somatomotor network showed the strongest amygdala dominance, while the visual network remained strongly hippocampus-dominant. Default mode and dorsal attention networks were moderately hippocampus-biased, and the frontoparietal network showed strong hippocampal dominance (noting that this was driven by relatively few parcels). The limbic network was near neutral or only weakly amygdala-biased, while the ventral attention network showed high variability with an overall mean close to zero (based on few parcels). Taken together, these results indicate that dominance and sharedness capture separable aspects of amygdalar- and hippocampal-cortical organization and relate to distinct axes.

### Dominance/sharedness relate to macroscale cortical organization, and sharedness is more estimator-sensitive (Figure 2 C-E)

We next examined whether the dominance and sharedness framework relates to broader principles of cortical organization and whether its main features are stable across GLASSO and Pearson connectivity estimators (**Fig. 2C-E**). Specifically, we related both measures to canonical FC gradients (G1, G2 and G3) of the cerebral cortex. G1 reflects the sensory-to-transmodal axis, G2 the sensorimotor/auditory-to-visual axis, and G3 an axis of internal-to-external information processing (Margulies et al., 2016). Overall, dominance and sharedness exhibited distinct associations with these macroscale gradients. Pearson-derived dominance showed its strongest positive association with G2 (Kendall τ = 0.42, *qMSR* < 0.05) and a negative association with G3 (τ = −0.33, *qMSR* < 0.01), but associations with G1 were not significant. Notably, because functional gradients are sign-ambiguous, these effects reflect associations with opposite poles of the same underlying organizational axes rather than directional effects per se. GLASSO-derived dominance showed a weaker but significant positive relationship with G2 (τ = 0.21, *qMSR* < 0.05).

Sharedness measures followed a distinct pattern. Pearson-derived sharedness was weakly positively associated with G1 (τ = 0.19, *qMSR* < 0.05) and negatively associated with G3 (τ = −0.42, *qMSR* < 0.01). GLASSO-derived sharedness showed a similar but more selective pattern, with a positive association with G1 (τ = 0.24, *qMSR* < 0.05), while associations with the remaining gradients were not statistically significant. Together, these patterns indicate that dominance and sharedness capture separable aspects of amygdalar-cortical and hippocampal-cortical organization and relate to distinct axes of macroscale cortical hierarchy, with partially convergent but non-redundant structure across connectivity estimators.

To test whether the count-based dominance metric reflected meaningful differences in connectivity magnitude rather than solely threshold-induced effects, we compared it against a complementary strength-based amygdala-minus-hippocampus contrast. Count-based dominance showed significant positive correspondence with the strength-based contrast in both estimators, with a stronger relationship observed for GLASSO (*r* = 0.62, *p_MSR_* < 0.001) than for Pearson (*r* = 0.43, *p_MSR_* < 0.05). These results indicate that the count-based formulation captures meaningful differences in relative connection strength rather than being driven by thresholding artefacts alone. We further recomputed the dominance and sharedness cortical maps with different mask thresholds. In both GLASSO and Pearson, the top 10% dominance and sharedness maps were highly corresponding with their top 5% and top 15% counterparts (**Supp. Fig. 1**).

Cross-estimator comparisons further revealed that dominance was more stable than sharedness. GLASSO- and Pearson-derived dominance maps were moderately well aligned (*τ* = 0.59, *p_MSR_* < 0.001), whereas sharedness exhibited substantially weaker agreement across estimators (*τ* = 0.25, *p_MSR_* < 0.01). These findings suggest that relative amygdala-hippocampus dominance constitutes a robust feature of cortical organization, while balanced convergence (sharedness) is more sensitive to whether connectivity is estimated from dense co-fluctuation structure (Pearson) or from sparser partial-correlation (GLASSO).

### Seed-specific cortical preference and intrinsic cross-structure coupling link local MTL interactions to distributed cortical convergence (Figure 3)

We next assessed how different hippocampal and amygdalar subregions preferentially targeted cortical territories with varying dominance or sharedness properties, and whether these extrinsic cortical preferences related to intrinsic cross-structure coupling between hippocampal and amygdalar subregions **(Fig. 3**).

**Figure 3.**
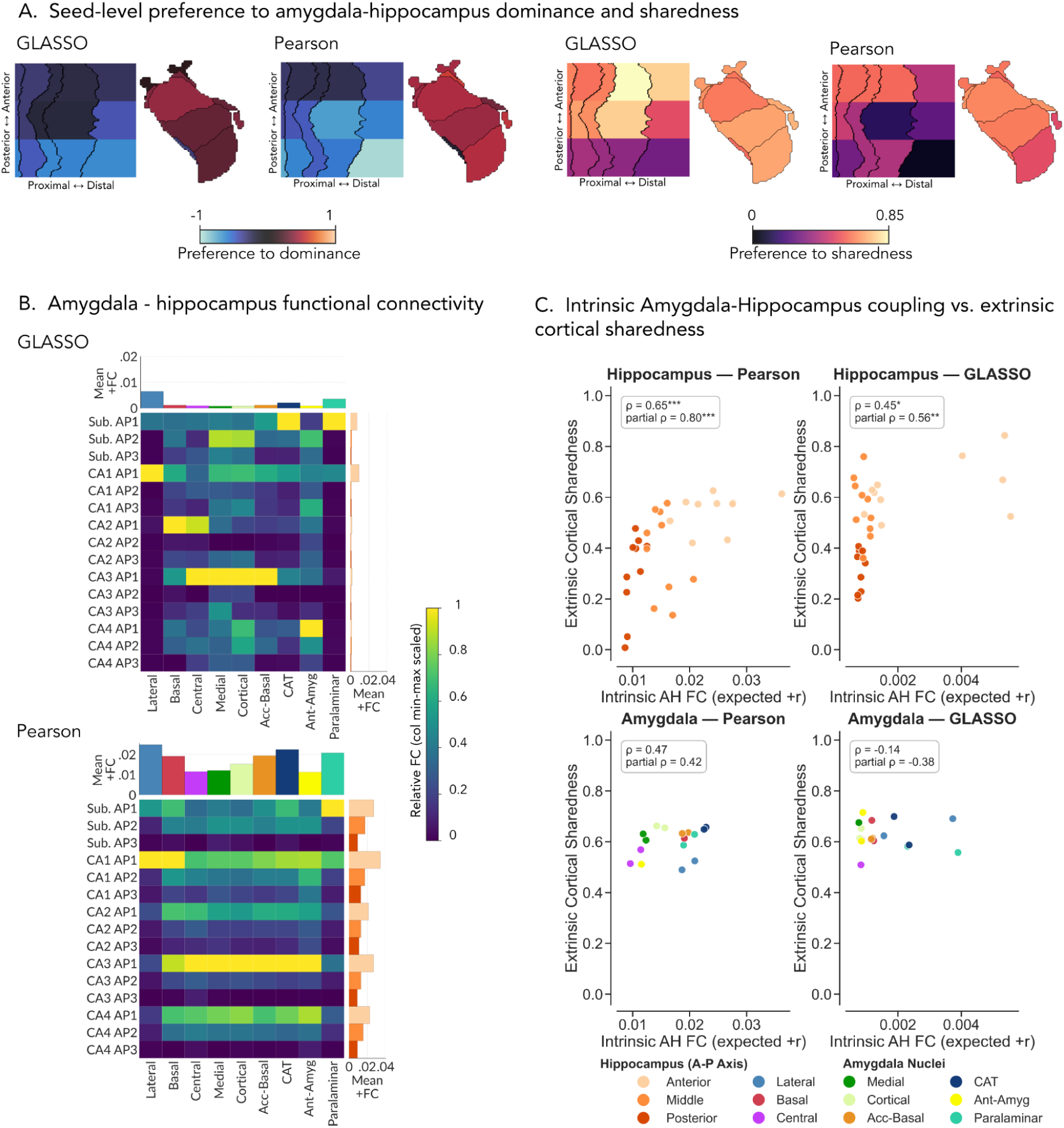
Seed-specific cortical preference and cross-structure seed-seed coupling. Only left hemisphere brain maps (and intrinsic coupling matrices) are shown for visualization purposes, analyses are conducted on the entire brain. Right hemisphere results can be found on **Supplementary Figure 3**. **A)** Hippocampal-cortical and amygdalar-cortical preferences were mapped onto hippocampal (top) and amygdalar subregions (bottom) using a Leave-One-Seed-Out (LOSO) strategy to reduce circularity. For each amygdala nucleus and hippocampal subregion, group-level preference to dominance (left) and sharedness (right) scores were additionally projected onto the corresponding subregions. Results are displayed separately for the GLASSO and Pearson estimators. For preference to dominance, positive values (warmer colors) indicate preferential weighting toward the amygdala-dominant cortex, whereas negative values (colder colors) toward the hippocampus-dominant cortex, and values near zero indicate balanced/weak preference. For preference to sharedness, higher values (lighter colors) indicate preferential weighting toward cortical parcels where both structures contribute strongly and in balanced contributions from both hippocampus and amygdala. **B)** Hippocampal-amygdalar FC between each hippocampal subfield and amygdalar nucleus pair is displayed as a heatmap, separately for GLASSO (top) and Pearson (bottom), ipsilateral shown only for simplicity). The color scale represents positive FC strength summarized as the unconditional positive mean across subjects (expected +r). Marginal bar plots show mean cross-structure coupling for each seed, and column-wise min-max scaled matrices highlight relative interaction structure independent of absolute strength, marginal bar colors represent different amygdala subnuclei and different hippocampus AP positions. **C)** Associations between extrinsic cortical sharedness and intrinsic cross-structure coupling at the seed level. Scatter plots illustrate the relationship between each seed’s intrinsic cross-structure coupling strength and its extrinsic cortical sharedness preference. Statistical annotations report both zero-order and partial Spearman correlations, with partial correlations controlling for the seed’s total positive FC strength. Asterisks indicate FDR-corrected significance (\**q* < 0.05, \*\**q* < 0.01, \*\*\**q* < 0.001).

Using a LOSO strategy to reduce circularity, we found that seed-specific preference to dominance was highly structure-consistent across both GLASSO and Pearson estimators. Hippocampal subregions exhibited negative dominance preference values, indicating preferential weighting toward hippocampus-dominant cortex, whereas amygdala nuclei showed positive values, reflecting preferential weighting toward amygdala-dominant cortex. Within the hippocampus, dominance preference varied systematically along the anterior-posterior (longitudinal) axis, with more posterior subregions showing stronger hippocampus-dominant preference and more anterior subregions showing weaker or more neutral values, thus relatively closer to amygdala-dominant cortical territory. Among amygdalar nuclei, the paralaminar nucleus was notable in showing comparatively neutral or hippocampus-leaning dominance preference across both estimators, standing out from the rest of amygdala.

In contrast to dominance, sharedness preference exhibited a more graded organizational pattern. Hippocampal subregions varied substantially in the extent to which they preferentially targeted highly shared cortical regions, mostly along the longitudinal axis, whereas amygdala subnuclei also showed variability, albeit over a narrower range. Thus, the cortex-level dominance and sharedness organization is recapitulated at the level of individual subregion embedding rather than being restricted solely to cortical parcels.

Intrinsic amygdala-hippocampus FC coupling also displayed pronounced subregion-specific structure rather than homogeneous organization. The amygdala-hippocampus FC matrices revealed clear row-and column-wise patterning across estimators. Prominent bands of increased coupling were observed particularly for anterior hippocampal subregions, indicating that cross-structure interactions were concentrated within specific hippocampal subregions rather than diffusely distributed across the matrix. Within the amygdala, lateral, paralaminar, and cortico-amygdaloid transition (CAT) nuclei showed especially selective relationship with hippocampal subregions: the lateral nucleus was preferentially coupled with anterior CA1, whereas paralaminar and CAT nuclei with anterior subiculum. Other amygdalar nuclei displayed broader connectivity patterns, although these interactions remained biased toward anterior hippocampal subregions. Column-wise normalized matrices further supported this selective organization by demonstrating strong relative selectivity, independent of absolute connectivity magnitude. These results indicate that local hippocampal-amygdalar coupling is highly structured at the subregional level.

We then tested whether intrinsic cross-structure coupling strength (using the mean positive connectivity of a subregion against the other structure) was associated with each subregion’s extrinsic preference for a shared cortical territory. This relationship proved to be asymmetric across structures. For hippocampal subregions, stronger intrinsic amygdala-hippocampus coupling predicted greater extrinsic cortical sharedness in both estimators (Pearson *ρ*=0.65, *q*<0.001; partial *ρ*=0.80, *q*<0.001 and GLASSO *ρ*=0.45, *q*<0.05; partial *ρ*=0.56, *q*<0.01). In contrast, analogous associations for amygdala subnuclei were non-significant in Pearson (*ρ* = 0.47, partial *ρ* = 0.42) and absent or negative in GLASSO (*ρ* = -0.14, partial *ρ* = -0.38). These findings suggest that hippocampal subregions exhibiting stronger local coupling with the amygdala also preferentially target cortical regions jointly embedded by both structures, while this relationship is not evident for amygdalar subnuclei.

Finally, to evaluate sampling stability, we repeated the masked map generation procedure across 200 random subject split-halves of the cohort. In each iteration, method-specific seed-wise top 10% masks, dominance and sharedness maps, seed-level LOSO preference scores, and amygdala-hippocampus coupling matrices were recomputed independently within each half. Both cortical dominance and sharedness maps demonstrated high split-half reliability. Pearson-derived maps approached ceiling stability, with median split-half correlations of *r* = 0.972 for dominance and *r* = 0.982 for sharedness. GLASSO-derived maps also showed strong stability despite their sparse support-prevalence formulation, with median split-half correlations of *r* = 0.873 for dominance and *r* = 0.953 for sharedness. Method-specific union masks were similarly stable for both estimators, with median Dice coefficients of 0.902 for GLASSO and 0.938 for Pearson. Seed-level LOSO sharedness scores were also highly reproducible across halves, with median correlations of *r* = 0.926 for GLASSO and *r* = 0.975 for Pearson. Finally, amygdala-hippocampus positive coupling matrices showed near-ceiling split-half reliability for both estimators, with median ipsilateral matrix correlations exceeding 0.994. Collectively, these analyses indicate that the observed cortical maps and seed-level organizational patterns were highly robust and not driven by idiosyncratic participant subsets.

### Joint gradients summarize the low-dimensional cortical embedding of hippocampal and amygdalar subregions (Figure 4)

Finally, we summarized the joint cortical embedding of hippocampal and amygdalar subregions in a lower-dimensional form by applying diffusion embedding to the full subregion-to-cortex FC matrices (**Fig. 4**). This approach enabled the identification of principal axes along which hippocampal and amygdalar subregions converge or diverge in their cortical connectivity profiles, thereby revealing whether these subregions segregate, interdigitate or vary along shared organizational gradients. The aligned hemispheric solutions revealed a robust low dimensional structure, with the first two gradients explaining 48.7% of the variance in the left hemisphere and 46.7% in the right hemisphere for GLASSO, and 49.9% in the left hemisphere and 51.1% in the right hemisphere for Pearson (**Supp. Fig. 2**).

**Figure 4.**
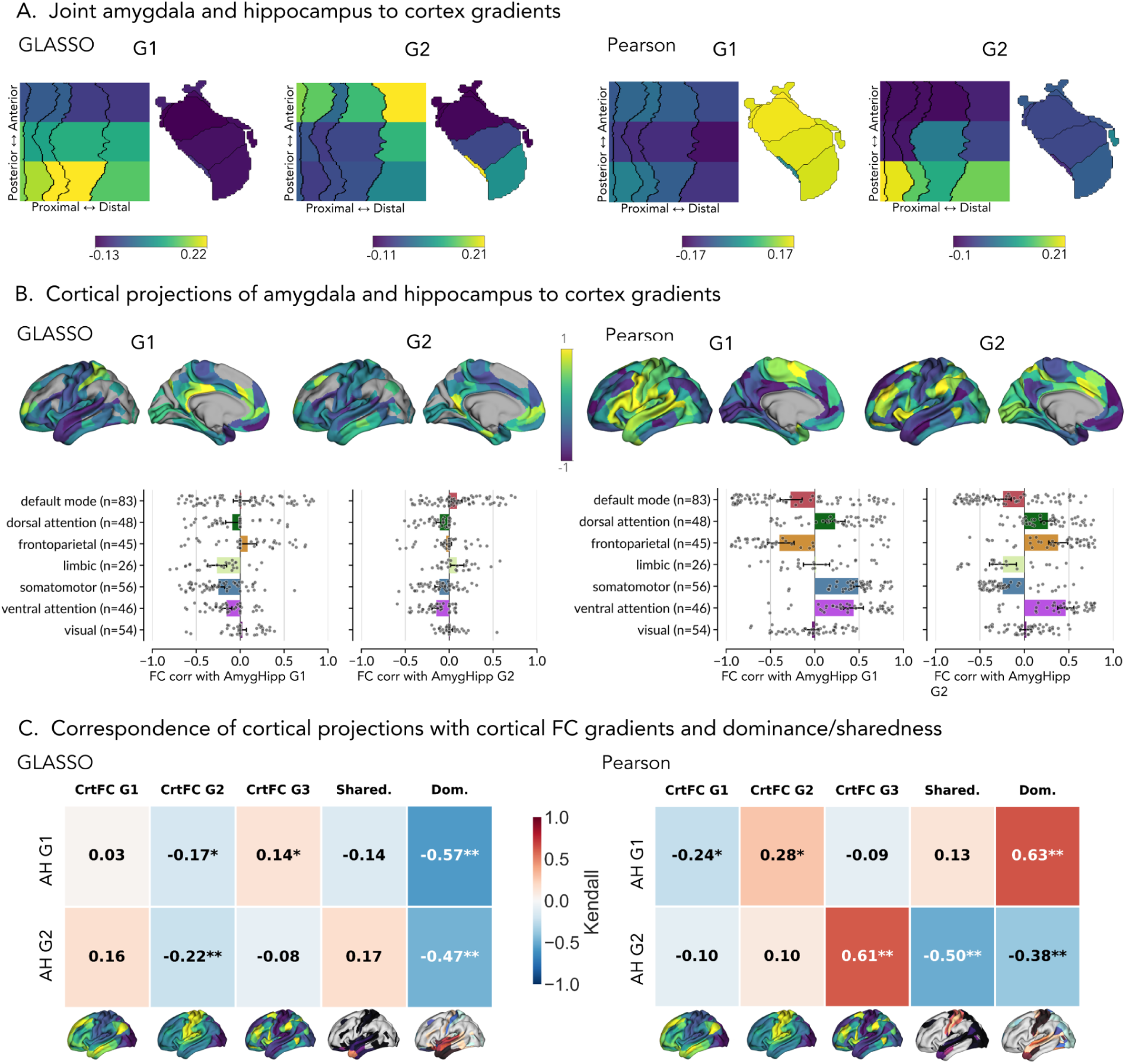
Joint amygdala and hippocampus to cortex gradients. Only left hemisphere brain maps are shown for visualization purposes, analyses are conducted on the entire brain. Right hemisphere results can be found on **Supplementary Figure 3**. All cortical surface maps are accompanied with plots showing the relevant values across Yeo-7 networks (bars = network means, lines = 95% CI, dots = parcels). **A)** Primary (G1) and secondary (G2) connectivity gradients of hippocampal-cortical (top) and amygdalar-cortical (bottom) FC. G1 and G2 are projected onto hippocampal and amygdalar subregions, separately for the GLASSO and Pearson estimators. Regions exhibiting similar colors reflect similar FC profiles, whereas divergent colors indicate greater dissimilarity in connectivity organization. **B)** Cortical projections of the joint hippocampal-cortical and amygdalar-cortical gradients (G1 and G2), estimated separately for the GLASSO (left) and Pearson (right) estimators. Higher projection values (light colors toward +1) indicate cortical parcels that are more strongly coupled with subregions with positive gradient scores (light colors in Panel A). Lower projection values (cool colors toward -1) indicate association with subregions located at negative ends of gradients (cool colors in Panel A). For each cortical projection, values are further summarized across the Yeo-7 networks. **C)** Heatmaps depict parcel-wise correspondence between cortical projections of joint hippocampal- and amygdalar-cortex gradients and canonical cortical functional gradients (CrtFC G1-G3), together with cortex-level amygdala-hippocampus dominance and sharedness maps. Results are shown separately for GLASSO and Pearson estimators. Asterisks indicate the significance of spatial correspondence assessed via Moran Spectral Randomization and FDR-corrected (* *q_MSR_*<0.05, \*\**q_MSR<_*0.01, *** *q_MSR_*<0.001).

The two estimators emphasized partially distinct aspects of this low-dimensional embedding. In the Pearson estimator, primary gradient (G1) captured a macroscale structure-level separation: amygdala nuclei occupied one end of the axis with relatively homogeneous high loadings (with the exception of the paralaminar nucleus, which aligned more closely with hippocampal profiles), while hippocampal subregions occupied the opposite end and exhibited additional internal variation. The secondary gradient (G2) reintroduced stronger within-hippocampus differentiation, especially along the anterior-posterior (longitudinal) axis with anterior hippocampal subregions showing greater similarity to amygdala, and more subtle differentiation across amygdalar nuclei.

In the GLASSO solution, amygdalar subregions appeared more homogenous along G1, while hippocampal subregions exhibited a pronounced internal gradient along the long axis, with anterior hippocampus positioned closer to amygdala and posterior hippocampus more distant; posterior CA1 emerged as the hippocampal subregion most distinct from amygdala. GLASSO G2 introduced additional differentiation within both hippocampus and amygdala, yielding a more interdigitated organization across the two structures. Within the amygdala, a paralaminar-to-basolateral-to-rest axis was evident, and within the hippocampus a subiculum to rest of hippocampus axis emerged, accompanied by an anterior to rest of hippocampus motif, with paralaminar nucleus and anterior subiculum occupying similar positions. Interestingly across the gradients there seemed to be a subtle within basolateral nucleus axis (most pronounced in GLASSO G2) that was ordered paralaminar, lateral, basal, accessory basal; with paralaminar nucleus being most similar to hippocampus across gradients.

When projected across the full retained cortical parcellation, these gradients yielded structured cortical maps that differed between estimators, reflecting differences between broad Pearson co-fluctuation and sparser GLASSO conditional association. In the GLASSO estimator, G1 and G2 showed relatively modest network-level separation, with limited variation across canonical functional networks. In contrast, the Pearson estimator exhibited stronger network differentiation: G1 contrasted frontoparietal and default mode networks against somatomotor, ventral attention and dorsal attention networks. G2 revealed an opposing axis positioning default mode, limbic and somatomotor networks at one end and frontoparietal, dorsal and ventral attention networks toward the other. Overall, these projections recapitulated key distinctions observed in the dominance and sharedness framework, while providing a continuous low-dimensional summary of subregion organization.

Correspondence analyses indicated that the cortical projections of the joint gradients were related both to canonical cortical functional gradients and to the top-10%-defined dominance/sharedness maps. For GLASSO, AmygHipp G1 showed significant correspondence with cortical gradient 3 (τ = 0.14, *qMSR* < 0.05), cortical gradient 2 (τ = -0.17, *qMSR* < 0.05), and cortical dominance (τ = -0.57, *qMSR* < 0.01). AmygHipp G2 showed significant correspondence with cortical gradient 2 (τ = -0.22, *qMSR* < 0.01) and cortical dominance (τ = -0.47, *qMSR* < 0.01). For Pearson, AmygHipp G1 showed significant correspondence with cortical gradient 2 (τ = 0.28, *qMSR* < 0.05), cortical dominance (τ = 0.63, *qMSR* < 0.01), and cortical gradient 1 (τ = -0.24, *qMSR* < 0.05). AmygHipp G2 showed significant correspondence with cortical gradient 3 (τ = 0.61, *qMSR* < 0.01), cortical sharedness (τ = -0.50, *qMSR* < 0.01), and cortical dominance (τ = -0.38, *qMSR* < 0.01). Because gradient polarity is sign-agnostic, positive and negative correspondence values should not be interpreted as intrinsically different effects. Rather, they indicate alignment with opposite ends of the displayed gradient axis. Together, these findings show that hippocampal and amygdalar subregions can be positioned within a shared low-dimensional cortical embedding that recapitulates both their segregated preference territories and their selective zones of convergence.

## Discussion

In this study, we examined hippocampal and amygdalar subregions within a shared cortical framework. Isocortical parcels did not separate into distinct hippocampal and amygdalar territories. Instead, they varied along a continuum from relative preference for one structure to more balanced contributions from both. This pattern was evident in both the dominance/sharedness maps and the joint gradients. Therefore, our findings extend systems-level accounts that have typically framed hippocampal organization with respect to anterior-temporal and posterior-medial cortical systems (Ranganath & Ritchey, 2012; Ritchey et al., 2015), and amygdala organization with respect to medial and orbitofrontal prefrontal, temporal, and paralimbic networks (Bickart et al., 2014; Aggleton et al., 2015). Across both FC estimators, this variation reflected either broader co-fluctuation patterns or associations conditioned on the rest of the network, and thus relatively less confounded by indirect relations.

The comparison between Pearson correlation and GLASSO emphasized different aspects of the same joint hippocampus-amygdala cortical embedding. Pearson produced broader cortical coverage, with stronger extension into visual, dorsal attention, and somatomotor territories. GLASSO yielded a sparser map that was concentrated more strongly towards limbic and ventral attention cortices, with notable overlap between methods in default mode and limbic regions. The frontoparietal cortex, by contrast, was largely outside both masks. That pattern aligned with the methodological distinction between the estimators: Pearson correlation captures dense co-fluctuation structure, including indirect and shared statistical dependencies, whereas regularized partial correlation conditions on the rest of the network and tends to yield a more focal estimate of conditional association (Friedman et al., 2008; Smith et al., 2011; Liégeois et al., 2020; Peterson et al., 2025). Interpreted together, these patterns suggest that Pearson and GLASSO do not simply provide stronger or weaker versions of the same map. Instead, they separate two organizational levels: a broad co-fluctuation field that may include indirect and shared-input dependencies, and a narrower conditional convergence core centered especially on paralimbic, medial temporal, and orbitofrontal areas.

Our dominance and sharedness analyses emphasize the areas across the cortex where the hippocampus or the amygdala was more repeatedly represented and where balanced co-representation emerges, rather than where on the cortex the two structures simply overlap. Because these maps were derived from the number of subregions from each structure that included a parcel within their top 10% cortical profile, they index relative representation rather than raw connectivity magnitude. From this perspective, the dominance maps identified cortical territories that were more consistently captured by one structure’s subregions than the other’s. This ordering was moderately stable across Pearson and GLASSO and positively corroborated (within parcels receiving contributions from both structures) by an independent strength-based amygdala-minus-hippocampus contrast. The strongest contrast in GLASSO dominance was between ventral attention, limbic and somatomotor networks on the amygdala dominant side against visual and default mode networks mainly on the hippocampus dominant side; while Pearson had somatomotor network as amygdala dominant and rest of the networks as hippocampus dominant (with limbic and ventral attention being neutral and also having a low number of parcels). Sharedness instead highlighted a smaller set of jointly represented cortical parcels and was more estimator-sensitive, with GLASSO emphasizing a focal limbic/paralimbic core, which is in line with primate anatomical work showing overlap between hippocampus and amygdala on the prefrontal cortex and medial temporal lobe (Saunders & Rosene, 1988a; Saunders et al., 1988b; Ghashghaei et al., 2007; Aggleton et al., 2015). Pearson yielded a pattern that extended more broadly, especially into the somatomotor cortex, compatible with human fMRI work utilizing pairwise correlation FC of amygdala and hippocampus with the isocortex (Ezama et al., 2021; Elvira et al., 2022; Klein-Flügge et al., 2022). Together, these findings suggest that the more robust feature of the joint embedding is not general hippocampus and amygdala overlap on the cortex, but a structured cortical landscape of relative representation with selective zones of balanced convergence. This interpretation is consistent with previous literature showing that cortico-hippocampal organization and amygdala subregional connectivity are both distributed across multiple large-scale systems rather than neatly confined to a single mnemonic or affective network (Ranganath & Ritchey, 2012; Sylvester et al., 2020; Elvira et al., 2022). Our results further indicate that different organizational scales can be captured using different FC methods.

At the subregion level, the clearest organizational structure was along the hippocampal long axis: posterior parcels were more closely associated with cortical areas that were more on the hippocampus dominant side, whereas anterior parcels shifted toward cortical territory that was less clearly hippocampal and more jointly represented by both structures. This was not only a dominance effect, as anterior hippocampal parcels also showed higher preference for shared cortical areas and stronger intrinsic amygdala coupling, suggesting that anterior hippocampus functions as the hippocampal interface with the amygdala-linked convergence zone. This pattern fits with long-axis accounts proposing that anterior hippocampus is more embedded in anterior-temporal, limbic, and medial-prefrontal circuits, whereas posterior hippocampus is more linked with posterior-medial/default mode network related systems (Ranganath & Ritchey, 2012; Poppenk et al., 2013; Strange et al., 2014). Overall, we show that the hippocampal long axis is expressed not only in relation to cortex, but also in how hippocampal subregions enter a joint cortical architecture shared with the amygdala. Within the amygdala, the repeated proximity of the paralaminar nucleus to the hippocampus was especially notable. That finding aligns with primate anatomical work showing that hippocampal inputs target paralaminar/basolateral territory and overlap immature neurons there, and more recent circuit work further showing that hippocampal pathways heavily innervate the intrinsic paralaminar basolateral nucleus, a zone linked to plasticity and context-sensitive processing (Fudge et al., 2012; Joyce et al., 2023). Human developmental studies likewise indicate that the paralaminar nucleus contains a large population of immature excitatory neurons that mature gradually across adolescence and persist into adulthood, making it a plausible biological substrate for the relatively hippocampus-like positioning observed here (Sorrells et al., 2019). The intrinsic-extrinsic association further underlined insights from cortical projections. Hippocampal subregions that were more strongly coupled to amygdala locally also tended to be linked to more jointly represented cortical territory, whereas the converse was not evident for amygdala nuclei. We interpret that asymmetry cautiously, but it is in keeping with the idea that some hippocampal subregions, especially more anterior and interface-like ones, may help link local amygdala coupling to distributed cortical integration of contextual and affective information, consistent with primate work on context-affect balance in amygdala microcircuits and recent human intracranial studies showing coordinated amygdala-hippocampus dynamics during emotional memory encoding and retrieval (Joyce et al., 2023; Qasim et al., 2023; Costa et al., 2025).

The joint gradients extended the dominance/sharedness analyses by showing that hippocampal and amygdalar subregions can be placed within a shared low dimensional space that preserves both structure-level separation and finer-grained interdigitation. In the Pearson estimator, the first gradient primarily behaved as a broad hippocampus-amygdala separation axis, whereas the second gradient reintroduced stronger within-hippocampus differentiation, especially along the long axis. In the GLASSO solution, the first gradient more strongly emphasized internal hippocampal variation, with anterior hippocampal parcels lying closer to amygdala, and the second gradient introduced additional within-amygdala heterogeneity, hippocampus subfield differentiation and cross-structure interdigitation. This pattern suggests that the joint embedding contains multiple organizational modes, some dominated by coarse structure-level differentiation and others by graded subregional similarity rather than a single clear-cut hippocampus-amygdala separation axis and that once broad indirect covariance is reduced, broader hippocampus-amygdala separation no longer occupies the dominant organization of the system, and finer subregional geometry becomes more visible. That interpretation fits well with recent work arguing that hippocampal-cortical organization itself is multidimensional rather than reducible to one long-axis gradient, including evidence for distinct anterior-posterior and secondary long-axis modes of cortical integration (Xie et al., 2024; Nordin et al., 2025). It also echoes human work indicating that amygdala subregions participate in distributed and partially dissociable cortical connectivity patterns, rather than forming a single uniform amygdala-to-cortex axis (Elvira et al., 2022; Sawada et al., 2022; Klein-Flügge et al., 2022). The cortical projections made these gradients interpretable as they aligned with canonical cortical gradients and with the count-based dominance/sharedness maps, indicating that the low-dimensional organization was anchored in recognizable macroscale cortical organization rather than arbitrary unexplainable dimensionality reduction.

One broader interpretation of these findings is that the cortical territories jointly represented by hippocampal and amygdalar subregions may mark sites where contextual-relational and affective/salience related signals are more likely to be coordinated, rather than implying a single undifferentiated medial temporal lobe system. This interpretation is in line with primate anatomical work showing that hippocampal and subgenual pathways in the amygdala are arranged in ways that may help balance context and affect processing, and with human intracranial studies showing coordinated amygdala-hippocampus dynamics during emotional memory encoding and subsequent retrieval-related reactivation (Joyce et al., 2023; Qasim et al., 2023; Costa et al., 2025). It is also consistent with contemporary developmental and evolutionary views of the amygdala as a heterogeneous mosaic with pallial and subpallial contributions rather than a unitary structure, which may help explain why some nuclei appear more hippocampus-like and others more distinct in their joint cortical embedding (Medina, 2023; Yu et al., 2023). From this perspective, the fact that paralaminar/basolateral territory repeatedly fell closer to hippocampal and shared cortical space than the rest of amygdala supports the idea that internal amygdala heterogeneity contributes to graded cross-structure organization (although it does not demonstrate this by itself). Framed this way, our results fit more naturally with distributed models of context-affect integration rather than with a strict separation between a memory system centered on hippocampus and an emotion system centered on amygdala.

Beyond this functional interpretation, our findings may also reflect deeper organizational principles rooted in cortical topography and evolutionary or developmental constraints. Based on dual origin theory (Dart, 1934; Pandya et al., 2015), cortical architecture has been described as reflecting partly dissociable ventral trends linked more closely to amygdala/piriform related (paleocortical) lineage and dorsal trends linked more closely to hippocampal (archicortical) lineage, with modern work extending this idea to connectivity and macrostructural organization in mammals and humans (Goulas et al., 2019; García-Cabezas et al., 2019; Valk et al., 2020). Our results do not test such developmental claims directly, but the repeated separation of more amygdala dominant and more hippocampus dominant cortical territory can be fitting with the idea that ancient ventral and dorsal organizational biases may still be visible in human systems-level functional embedding. However, our data do not suggest two clearly separated streams. Instead, the coexistence of relative dominance, selective sharedness, and cross-structure interdigitation in the joint gradients may fit better with recent parallel-systems models showing that association cortex contains multiple closely juxtaposed distributed networks with distinct medial temporal affiliations, including separable medial temporal lobe linked default mode network and social/association network branches (Braga & Buckner, 2017; DiNicola et al., 2020; Edmonds et al., 2024; Reznik et al., 2023, 2024; Braga, 2025). Based on this perspective, the present findings cannot be viewed as evidence for two pure developmental systems by themselves, but as a subregion-level functional mapping of how hippocampal and amygdalar subregions participate in partially parallel and partially convergent cortical patterns that may echo ontogenetic ventral-dorsal organizational principles.

A final set of limitations should temper interpretation. First, our findings are based on resting-state FC and therefore do not capture how hippocampal and amygdalar subregions engage the cortex during behavior, emotional memory formation, or naturalistic cognition. Second, dominance and sharedness maps are count-based summaries of relative top 10% subregional representation, not direct measures of connectivity strength or causal influence, though the dominance maps were positively corroborated by an independent strength-based amygdala-minus-hippocampus contrast. Importantly, because these maps are derived from each subregion’s own relative connectivity profile, a subregion with weak overall connectivity can still contribute to a parcel if that parcel ranks in its top 10%, meaning these maps reflect *relative cortical representation across subregions*, not absolute coupling strength. Third, while GLASSO provides a sparser, conditionally specific estimate compared to Pearson correlation, it remains a statistical model of FC and does not capture effective or structural connectivity (Smith et al., 2011; Liégeois et al., 2020; Peterson et al., 2025). A related but distinct limitation concerns signal quality: the subcortical signal, particularly in the hippocampus and amygdala, inherently has a low signal-to-noise ratio (SNR) in standard resting-state fMRI, and the HCP-YA data were not specifically optimized for subcortical resolution or denoising. This may limit sensitivity to subtle subregional differences and could bias connectivity estimates, especially in smaller or more metabolically active nuclei. Finally, the two estimators were not designed for direct comparison: GLASSO masks were defined as the top 10% most prevalent positive connections per subregion (to handle sparsity), whereas Pearson masks used the top 10% strongest connections. As such, the results should be viewed as complementary rather than competing. Future work should test whether the same cortical preference, convergence, and gradient axes reconfigure during emotional or memory tasks, and whether this joint embedding is altered in neuropsychiatric conditions involving hippocampal or amygdalar dysfunction.

## Data availability

This study used publicly available neuroimaging data from the Human Connectome Project Young Adult dataset. These data are available to approved users through the Human Connectome Project data access platform: https://www.humanconnectome.org/study/hcp-young-adult/document/hcp-young-adult-2025-release and are subject to the relevant data use terms and restrictions. No new raw neuroimaging data were generated for this study.

## Code availability

Code used to run the analyses and generate figures is available at: https://github.com/CNG-LAB/amygdala_hippocampus_cortical_convergence_divergence. Raw Human Connectome Project data are not redistributed in the repository.

## Funding

DYE was funded by Max Planck Society. M.M. was funded by the Jacobs Foundation Research Fellowship. A.J. was funded by Max Planck Society. JdK is supported by a Natural Sciences and Engineering Research Council of Canada - Postdoctoral Fellowship (NSERC-PDF). JR was funded by Banting Fellowship. BCB acknowledges research support from the National Science and Engineering Research Council of Canada (NSERC RGPIN- 2025- 05932), CIHR (FDN- 154298, PJT- 174995, PJT-191853, PJT- 203761), SickKids Foundation (NI17- 039), HIBALL, Healthy Brains and Healthy Lives (HBHL), Brain Canada Foundation, FRQS, the Tier- 2 Canada Research Chairs Program, and the Centre for Excellence in Epilepsy at the Neuro (CEEN). SLV was funded by the Max Planck Society through the Lise Meitner Excellence Program, the Jacobs Foundation Research Fellowship, the Hector Research Career Development Award, the European Research Council (ERC) Starting Grant, SOCO.

## Role of the funders

The funders supporting the authors had no role in the design of the present study, data analysis, interpretation of the results, preparation of the manuscript, or the decision to submit the work for publication.

## Acknowledgements

Data were provided by the Human Connectome Project, WU-Minn Consortium (Principal Investigators: David Van Essen and Kamil Ugurbil; 1U54MH091657) funded by the 16 NIH Institutes and Centers that support the NIH Blueprint for Neuroscience Research; and by the McDonnell Center for Systems Neuroscience at Washington University.

## Ethics and Consent

This study involved secondary analysis of data from the Human Connectome Project Young Adult dataset. Data collection for the Human Connectome Project was approved by the relevant institutional review board at Washington University in St. Louis, and all participants provided written informed consent. The present study involved no new participant recruitment, intervention, or contact and used data accessed in accordance with the Human Connectome Project data-use terms.

## Competing interests

JDK and BCB are co-founders of BrainScores and hold stock.

## Author contributions

Conceptualization: Doruk Yiğit Erigüç, Sofie L. Valk.

Methodology: Doruk Yiğit Erigüç, Mylla Marsiglia, Alexandra John, Şeyma Bayrak, Jordan DeKraker, Sofie L. Valk.

Software: Doruk Yiğit Erigüç. Validation: Doruk Yiğit Erigüç.

Resources: Doruk Yiğit Erigüç, Bin Wan, Sofie L. Valk.

Investigation: Doruk Yiğit Erigüç, Anton Jakovčić.

Data curation: Doruk Yiğit Erigüç. Formal analysis: Doruk Yiğit Erigüç.

Visualization: Doruk Yiğit Erigüç, Şeyma Bayrak. Writing—original draft: Doruk Yiğit Erigüç.

Writing—review and editing: Doruk Yiğit Erigüç, Mylla Marsiglia, Alexandra John, Şeyma Bayrak, Bin Wan, Anton Jakovčić, Jessica Royer, Boris C. Bernhardt, Sofie L. Valk.

Project administration: Doruk Yiğit Erigüç, Sofie L. Valk. Supervision: Sofie L. Valk.

Funding acquisition: Sofie L. Valk.

## Notes

https://www.humanconnectome.org/study/hcp-young-adult

### Summary of Updates

Added author contributions, ethics; updated participants section; changed github link; clarified that cortical gradients are sample-derived; typos corrected.

