## Supplemental Figures for "Selective convergence and graded divergence of hippocampal and amygdala subregions using functional connectivity"

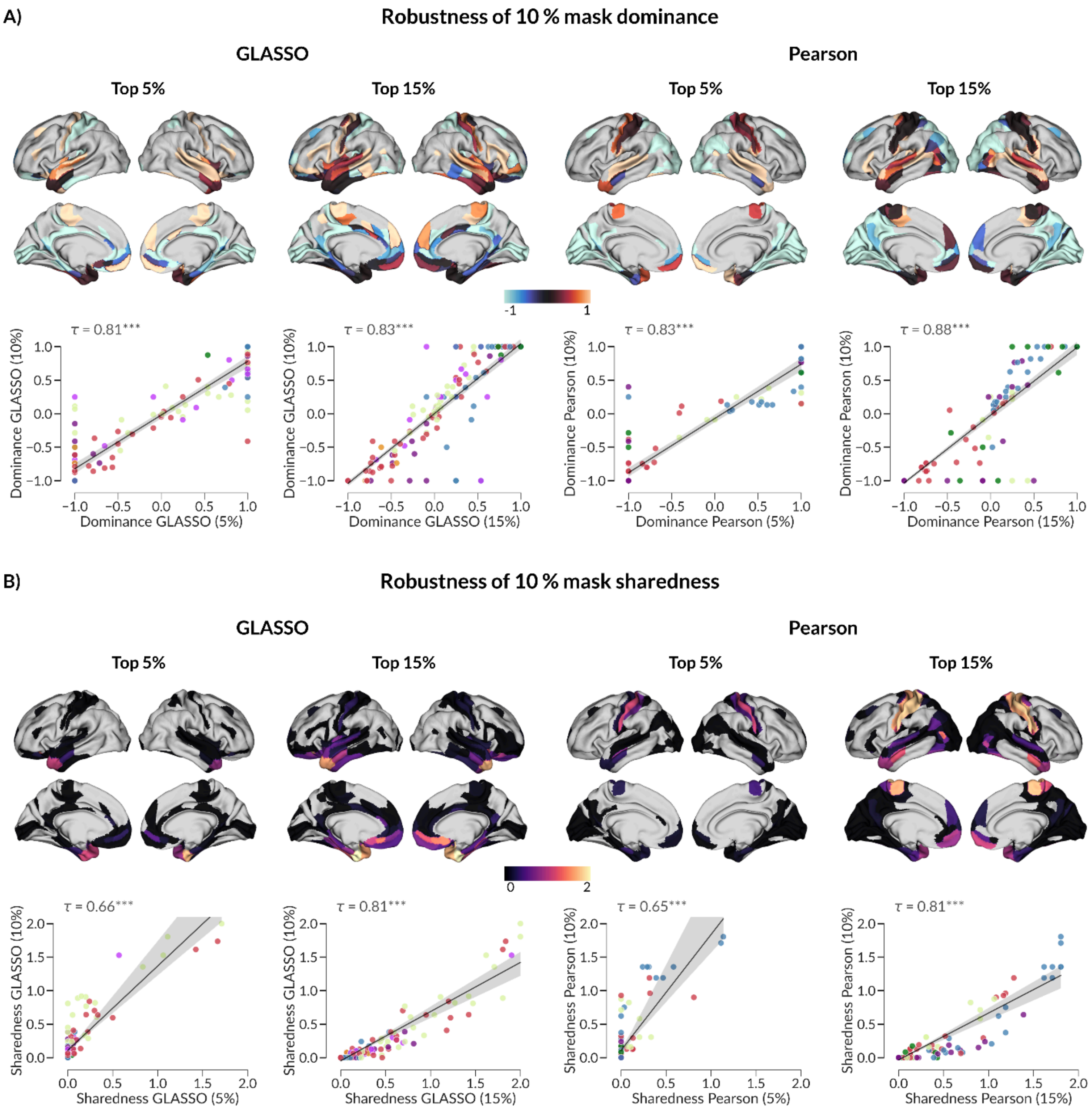


**Supplementary Figure 1. Dominance and sharedness metrics are robust to different threshold masks.** Dominance **(A)** and sharedness **(B)** maps derived at 5% and 15% thresholds showed high correspondence with the 10% maps for both Pearson and GLASSO estimates. Kendall correlations are shown for descriptive robustness assessment. Asterisks indicate the significance of spatial correspondence assessed via Moran Spectral Randomization (* $p_{MSR}$<0.05, **$p_{MSR<}$0.01, *** $p_{MSR}$<0.001).

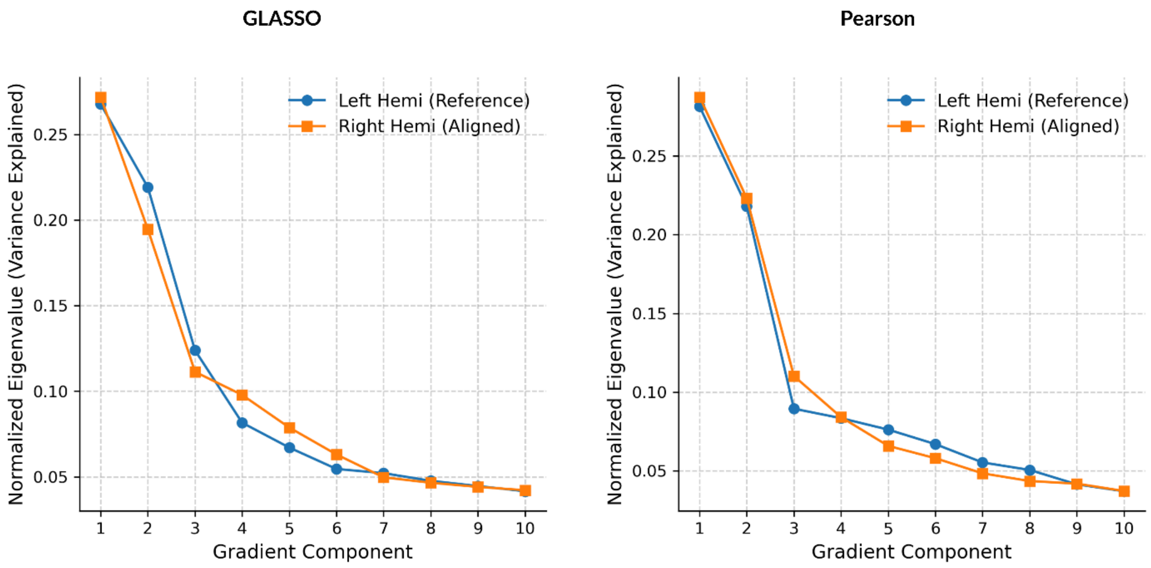


**Supplementary Figure 2. Variance explained by the gradient solutions across GLASSO and Pearson.** Normalized eigenvalues are shown for the first 10 gradient components derived from the left hemisphere and aligned right hemisphere solutions for GLASSO (left) and Pearson correlation (right).

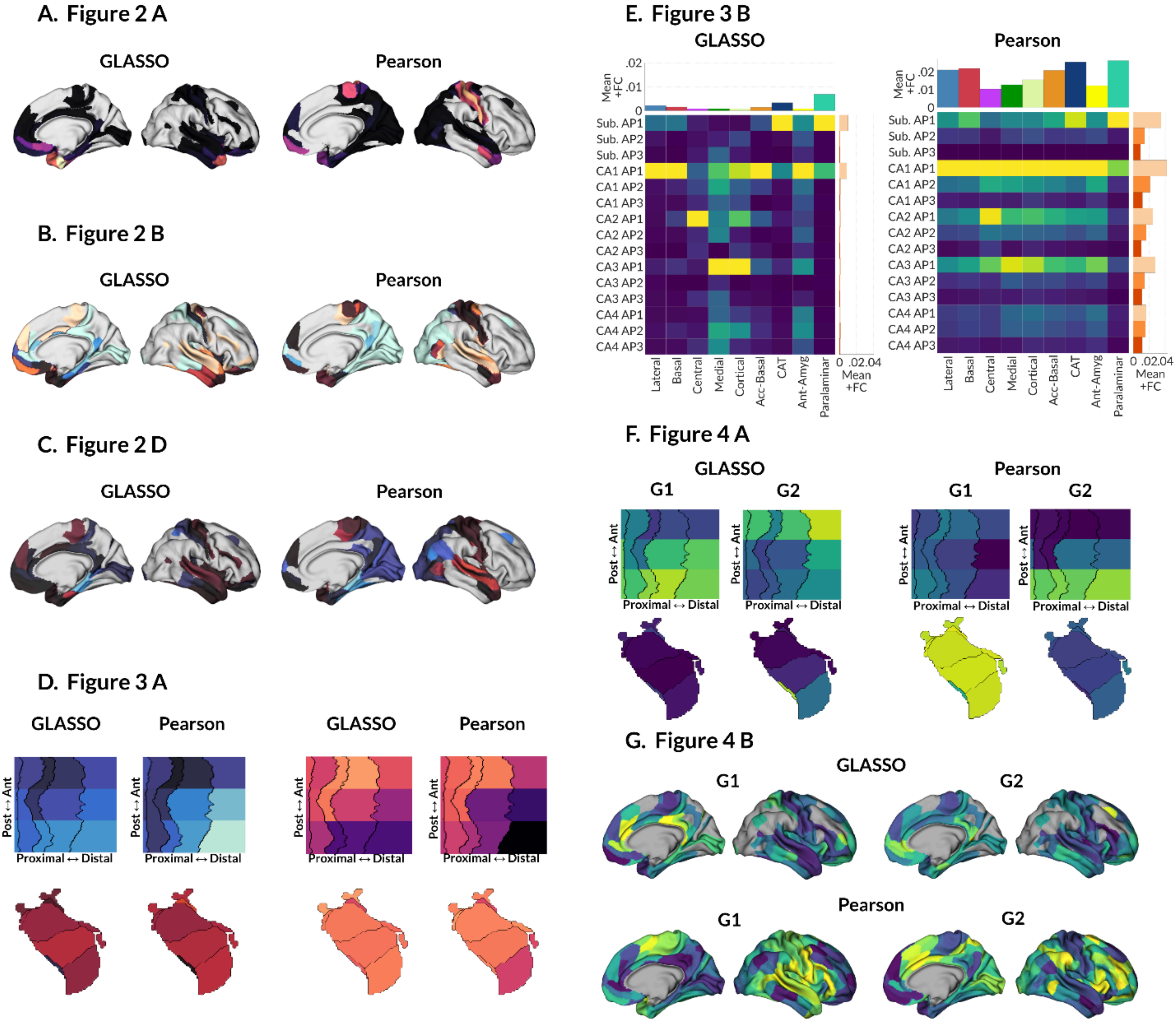


**Supplementary Figure 3. Right hemisphere counterparts of results from various figures.** More information and related colorbars can be found on the respective figures. **A)** Sharedness maps from **Figure 2 A**. **B)** Dominance maps from **Figure 2 B**. **C)** Strength based dominance maps from **Figure 2 D**. **D)** Seed-level preference to dominance (left) and sharedness (right) from **Figure 3 A**. **E)** Intrinsic amygdala-hippocampus FC matrices from **Figure 3 B**. **F)** Joint amygdala-hippocampus-to-cortex gradients from **Figure 4 A**. **G)** Cortical projections of gradients from **Figure 4 B**.
